# How do packet losses affect measures of averaged neural signals?

**DOI:** 10.1101/2021.06.03.446830

**Authors:** Evan M. Dastin-van Rijn, Matthew T. Harrison, David A. Borton

## Abstract

Recent advances in implanted device development have enabled chronic streaming of neural data to external devices allowing for long timescale, naturalistic recordings. However, characteristic data losses occur during wireless transmission. Estimates for the duration of these losses are typically uncertain reducing signal quality and impeding analyses. To characterize the effect of these losses on recovery of averaged neural signals, we simulated neural time series data for a typical event-related potential (ERP) experiment. We investigated how the signal duration and the degree of timing uncertainty affected the offset of the ERP, its duration in time, its amplitude, and the ability to resolve small differences corresponding to different task conditions. Simulations showed that long timescale signals were generally robust to the effects of packet losses apart from timing offsets while short timescale signals were significantly delocalized and attenuated. These results provide clarity on the types of signals that can be resolved using these datasets and provide clarity on the restrictions imposed by data losses on typical analyses.

## I. Introduction

In order to aid in the development of closed-loop therapies, many implanted device manufacturers have designed “bidirectional” implants capable of concurrently stimulating and sensing [1]–[4]. Early bidirectional devices stored data locally on the implant or required restrictive interfaces for wireless transmission of data limiting recordings to short time periods and unnatural environments. Furthermore, extensive sensing ran the risk of premature battery failure shortening device lifetime [5]. Recent devices such as the Medtronic Summit RC+S solve both limitations via rechargeable capabilities and improvements enabling chronic streaming of neural data to external devices up to 12 meters away. These advances allow access to long timescale neural recordings in natural environments facilitating the identification and development of personalized biomarkers and therapies [6]–[8].

During wireless transmission, samples of neural data are grouped into formatted units called “packets” [9]. Packets contain a series of subsequent samples of a particular length as well as timing information and other relevant metadata. When transmitted, it is possible for packets to fail to reach the receiver leading to missing samples. These missing samples need to be properly accounted for when the time series is reconstructed. The timing information contained in each packet aids in this process but is frequently inexact resulting in uncertainty in the number and location of the samples missing from a recording. This process, known as packet loss, is illustrated in Fig. 1.

**Fig. 1.**
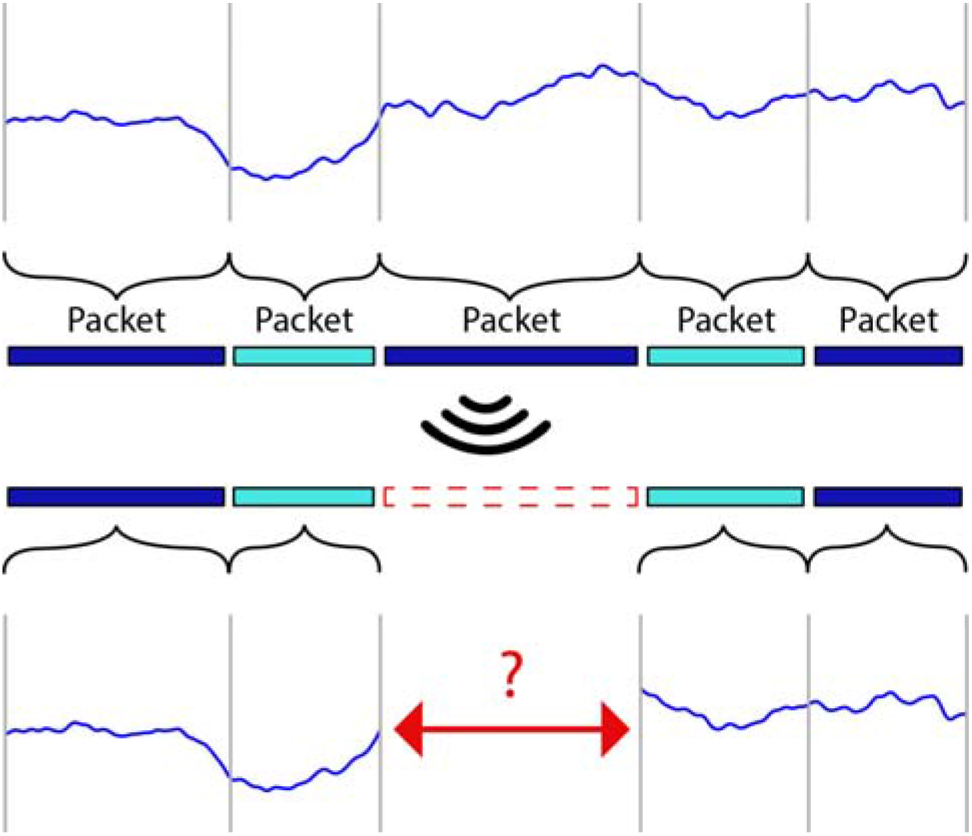
Illustration of packet loss. Subsequent samples from a neural data time series acquired on an implanted device are grouped into packets. Packets are then wirelessly transmitted to a receiver. During the transmission process it is possible for some packets to be lost. As a result, the relative timing of the samples contained in received packets is uncertain.

Uncertainty in packet timing has implications for recovering biological signals since the estimated time for each sample may be somewhat offset from the true time when the sample was recorded. We hypothesized that these effects would likely compound when trials were averaged leading to timing offsets, delocalization, reduction in amplitude, and difficulty resolving small differences between signals. Such changes would impact any analyses based upon evoked response potentials, peristimulus time histograms, or event related potentials. In this work, we sought to determine how the degree of timing uncertainty following a loss could influence the ability to accurately identify features of a typical averaged signal for which we simulated event-related potentials (ERPs) of variable duration.

## II. Methods

### A. ERP Simulation

In order to empirically determine the effects of packet losses on neural data, we simulated a positive-going ERP peak to provide a ground truth signal of interest. Each ERP trial ranged from −500 to 1500 ms and was modeled using a Gaussian adjusted to have an amplitude of 1 μV and a latency of 750 ms. We considered a range of standard deviations from 10-200 ms in order to approximate ERPs of variable duration [10]. All the simulated ERPs are shown in Fig. 2A.

**Fig. 2.**
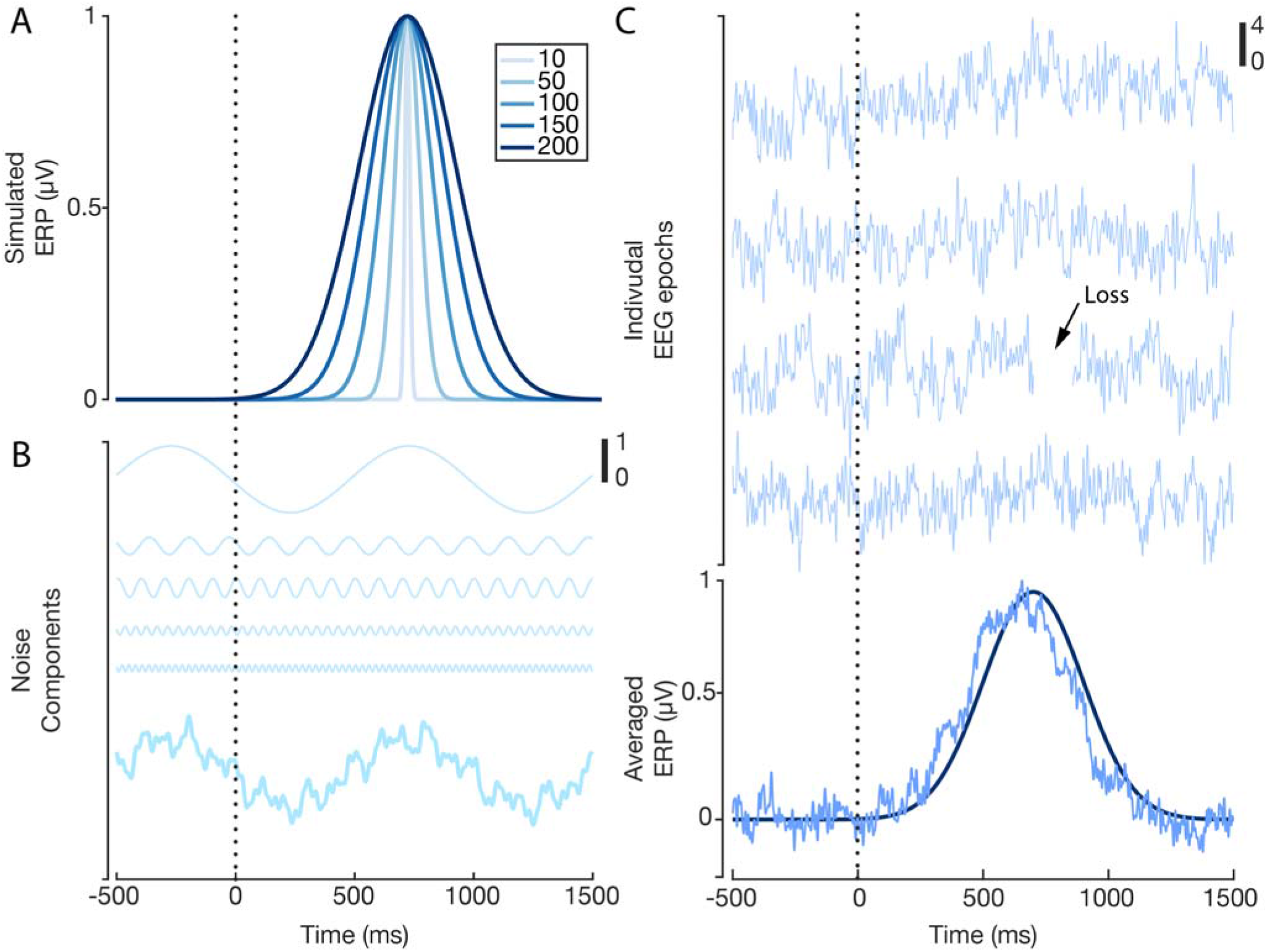
Illustration of ERP and noise simulations. (A) Ground-truth simulated Gaussian ERP trials for five different durations of interest. (B) Visualization of sinusoidal noise components. (C) Example ERP trials with noise and losses. Averaged signal is shown along with the ground-truth ERP in dark blue.

Each ERP trial was combined with simulated pink noise that was generated by summing together 50 sinusoids with randomly varying frequency and phase (with different random values for each simulated trial) [11]. Frequencies spanned the range from 0.1-125 Hz and the phase randomly varied between 0 and 2*π*. The maximum amplitude of any single frequency component of the noise was set to be 1 μV with the relative amplitude of each component scaled to match the power spectrum of typical electroencephalography data. The noise time series was divided by its standard deviation prior to being added to the ERP in order to ensure a signal to noise ratio of 20dB. Example sinusoids and a noise timeseries are shown in Fig. 2B. A total of 300 trials with a sampling rate of 1000 Hz were simulated for each experiment and concatenated into a single timeseries. This timeseries was then divided into 100 ms packets of which 30 were randomly removed, corresponding to 0.5% data loss. The estimated size of each loss was then randomly varied according to a normal distribution with a mean of 100 ms and a standard deviation ranging from 0 to 50 ms in order to consider a range of potential timing uncertainties. Packets were then recombined into a continuous time series with missing samples represented using NaNs. Lastly, the timeseries was divided into 300 trials and averaged to estimate the ERP. A typical example of this process is shown in Fig. 2C.

### B. ERP Measures

In order to compare the ERPs in the presence of packet losses to the ground truth signal, we looked at five different metrics: timing offset, delocalization, amplitude ratio, statistical significance, and effect size. To compute the first three metrics, we fit a Gaussian to the averaged signal by minimizing their mean squared error using the Matlab (Mathworks, Natick, MA, USA) function *fminsearch*. We defined timing offset as the difference in the mean between the fit and the ground truth signal (Fig. 3A), delocalization as the difference in the standard deviation between the fit and the ground truth signal (Fig. 3B), and the amplitude ratio as the amplitude of the fit signal divided by the amplitude of the ground truth signal (Fig. 3C). These metrics were chosen in order to determine how packet losses might shift, spread, and reduce estimates of biological signals. Additionally, we divided ERP trials into two conditions, where the second had ¾ of the amplitude of the first to estimate the effect of packet losses on discerning small differences between experimental conditions. For each trial, we computed the average amplitude within half a standard deviation of 750 ms and combined these measures to develop two a distribution for each condition. These distributions were compared on the basis of a two-sample t-test and Cohen’s d effect size.

**Fig. 3.**
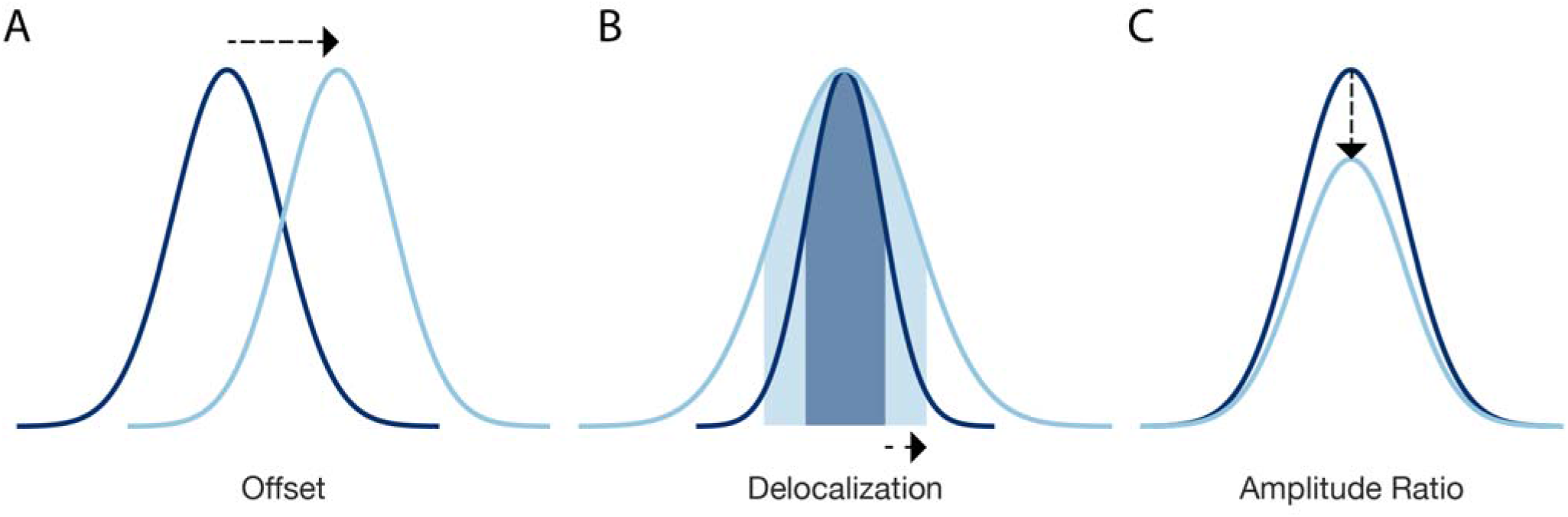
Illustration of ERP comparison metrics. (A) Offset describes the difference in mean between the recovered and ground-truth signals. (B) Delocalization describes the change in standard deviation between the recovered and ground truth signals. (C) Amplitude ratio is the ratio of the recovered signal amplitude to the ground-truth amplitude.

## III. Results

ERP comparisons were simulated and represented using heatmaps for signal durations ranging from 10-200 ms and timing errors ranging from 0-50 ms. Offset generally increased for larger timing errors with the greatest effect for signal durations above 50 ms. Offsets reached 50 ms for 25 ms timing error and 100 ms for 50 ms timing error (Fig. 4A). Delocalization had the opposite trend with respect to duration showing large increases for lower signal durations but a similar increase with respect to timing error. Delocalization reached 10 ms for longer signals and 100 ms for shorter signals (Fig. 4B). Amplitude ratio decreased for increasing timing error with greater and more rapid decreases for lower signal duration. Reduction in amplitude was minimal for 200 ms signals, 80 ms signals were reduced by 50% for timing errors greater than 25 ms, and 10 ms signals were completely attenuated for timing errors greater than 20 ms (Fig. 4C). significance probability (Fig. 4D) and effect size (Fig. 4E) showed similar trends to amplitude ratio with minimal effects for long signals, reduction in ability to resolve slight differences for 80 ms signals with 25 ms of timing error, and complete inability to resolve signal differences for short duration signals for increasing timing error.

**Fig. 4.**
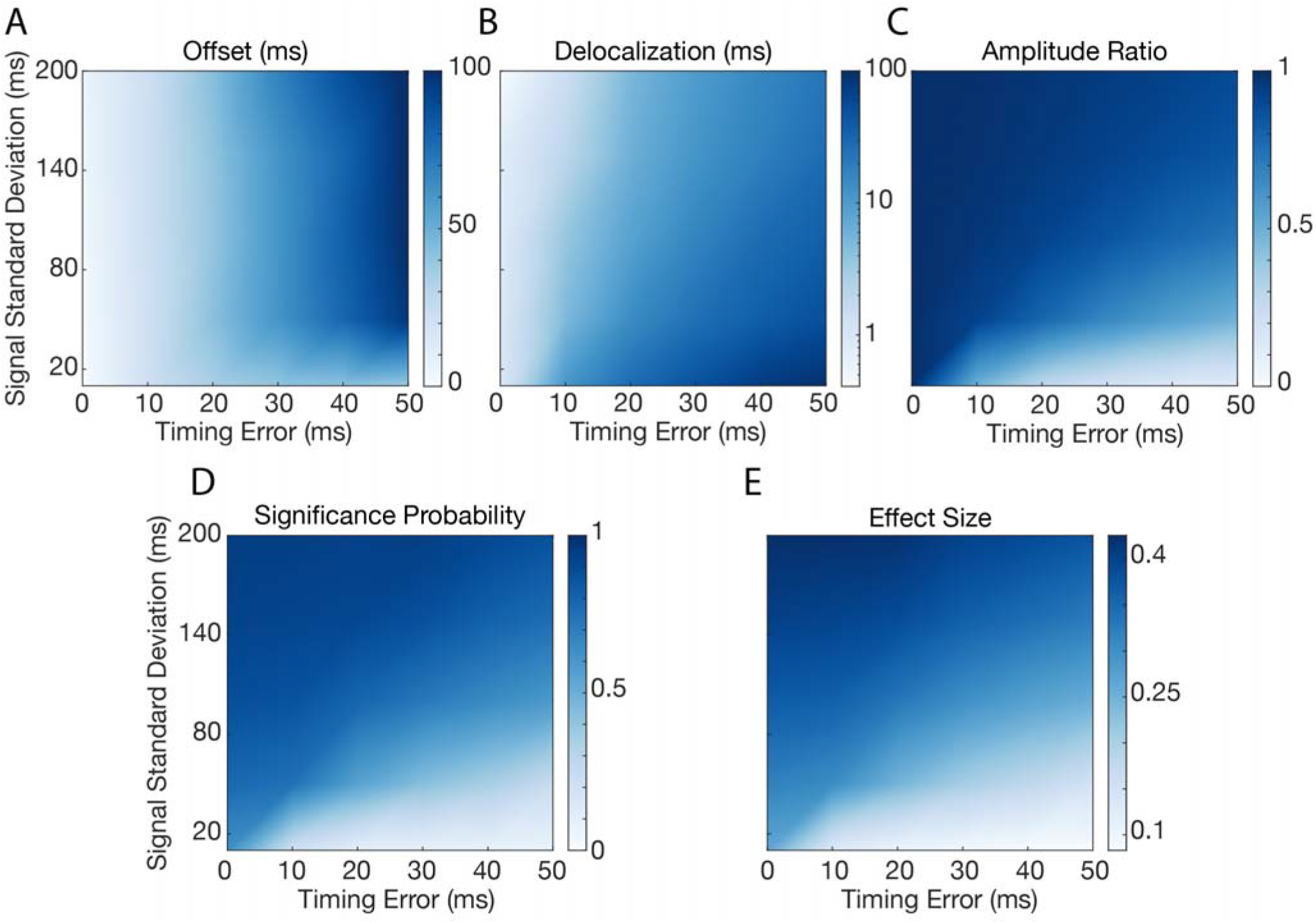
Simulation results. Heatmaps display the comparisons between the recovered ERP and ground-truth ERP as a function of the signal duration and the timing error when estimating losses for (A) offset, (B), delocalization, (C) amplitude ratio, (D) the proportion of trials which were significant, and (E) effect size.

## IV. Discussion

The aim of this work was to characterize the effect of packet losses on analyses of averaged signals for which we considered event related potentials depending on the signal duration and the degree of uncertainty in estimating the loss size. Our results indicate that packet losses have noticeable effects on averaged signals that lead to offsets in time, delocalization in time, reductions in amplitude, and increased difficulty resolving small differences between signals. These effects are generally greatest for shorter duration signals and for larger degrees of timing uncertainty with the exception of timing offsets. Offsets were largely identical across signal durations apart from particularly short signals which were effectively eliminated completely for higher timing uncertainties. Delocalizing effects were minimal for higher duration signals but severely impacted short signals for greater degrees of timing uncertainty. Amplitude ratio, statistical significance, and effect size were largely correlated with noticeable reductions occurring for signals with standard deviations below 80 ms.

For the purposes of this simulation, we chose to use a constant loss size of 100 ms and 30 losses in total. These values were chosen to be consistent with our experience using the Medtronic Summit RC+S with recordings in the clinic. However, our observations are relevant to any application where packetized data transmission is a requirement including implanted devices like the Summit RC+S [1], Active PC+S [12], and Neuropace RNS [8], wireless EEG systems like the B-Alart, Enobio, Muse, and Mindwave [13], and wireless EMG systems like the Athos [14] and Delsys Trigno [15]. In each device, characterizing the effects of data loss on signal averages is fundamental to ensuring accurate and effective analyses.

## V. Conclusion

Our results demonstrate that long timescale signals with standard deviations greater than 100 ms are robust to the effects of packet losses on averaged timeseries data. However, the exact timing of the signal onset will be uncertain to the level of twice the estimated timing uncertainty. Shorter timescale signals with standard deviations less than 80 ms are noticeably obscured by packet losses. Future studies interested in short timescale effects will need to be wary of potential spurious results due to packet losses or ensure sufficiently low timing uncertainty to avoid these effects all together. We hope that these results will help inform any future studies relying on signal averages in the presence of uncertain data losses.

## References

[1] S. Stanslaski et al., “A Chronically Implantable Neural Coprocessor for Investigating the Treatment of Neurological Disorders,” IEEE Trans. Biomed. Circuits Syst., vol. 12, no. 6, pp. 1230–1245, Dec. 2018.

[2] F. T. Sun and M. J. Morrell, “The RNS System: responsive cortical stimulation for the treatment of refractory partial epilepsy,” Expert Rev. Med. Devices, vol. 11, no. 6, pp. 563–572, Nov. 2014.

[3] S. Stanslaski et al., “Design and validation of a fully implantable, chronic, closed-loop neuromodulation device with concurrent sensing and stimulation,” IEEE Trans. Neural Syst. Rehabil. Eng., vol. 20, no. 4, pp. 410–421, Jul. 2012.

[4] A. Goyal et al., “The development of an implantable deep brain stimulation device with simultaneous chronic electrophysiological recording and stimulation in humans,” Biosens. Bioelectron., vol. 176, p. 112888, Dec. 2020.

[5] N. C. Swann et al., “Chronic multisite brain recordings from a totally implantable bidirectional neural interface: experience in 5 patients with Parkinson’s disease,” J. Neurosurg., vol. 128, no. 2, pp. 605–616, Apr. 2017.

[6] R. Gilron et al., “Chronic wireless streaming of invasive neural recordings at home for circuit discovery and adaptive stimulation,” bioRxiv, bioRxiv, 14-Feb-2020.

[7] V. Kremen et al., “Integrating Brain Implants With Local and Distributed Computing Devices: A Next Generation Epilepsy Management System,” IEEE J Transl Eng Health Med, vol. 6, p. 2500112, Sep. 2018.

[8] Z. M. Aghajan et al., “Theta oscillations in the human medial temporal lobe during real-world ambulatory movement,” Curr. Biol., vol. 27, no. 24, pp. 3743–3751.e3, Dec. 2017.

[9] K. Bazaka and M. V. Jacob, “Implantable Devices: Issues and Challenges,” Electronics, vol. 2, no. 1, pp. 1–34, Dec. 2012.

[10] P. E. Clayson, S. A. Baldwin, and M. J. Larson, “How does noise affect amplitude and latency measurement of event-related potentials (ERPs)? A methodological critique and simulation study,” Psychophysiology, vol. 50, no. 2, pp. 174–186, Feb. 2013.

[11] N. Yeung, R. Bogacz, C. B. Holroyd, and J. D. Cohen, “Detection of synchronized oscillations in the electroencephalogram: An evaluation of methods,” Psychophysiology, vol. 41, no. 6, pp. 822–832, Nov. 2004.

[12] E. G. M. Pels et al., “Stability of a chronic implanted brain-computer interface in late-stage amyotrophic lateral sclerosis,” Clin. Neurophysiol., vol. 130, no. 10, pp. 1798–1803, Oct. 2019.

[13] E. Ratti, S. Waninger, C. Berka, G. Ruffini, and A. Verma, “Comparison of medical and consumer wireless EEG systems for use in clinical trials,” Front. Hum. Neurosci., vol. 11, p. 398, Aug. 2017.

[14] S. K. Lynn, C. M. Watkins, M. A. Wong, K. Balfany, and D. F. Feeney, “Validity and reliability of surface electromyography measurements from a wearable athlete performance system,” J. Sports Sci. Med., vol. 17, no. 2, pp. 205–215, Jun. 2018.

[15] I. Poitras, M. Bielmann, A. Campeau-Lecours, C. Mercier, L. J. Bouyer, and J.-S. Roy, “Validity of wearable sensors at the shoulder joint: Combining wireless electromyography sensors and inertial measurement units to perform physical workplace assessments,” Sensors (Basel), vol. 19, no. 8, p. 1885, Apr. 2019.

